# *Uromyces plumbarius* first reported on *Oenothera lindheimeri* in Europe with widespread distribution and association with two common natural enemies of rust fungi

**DOI:** 10.1101/2025.04.07.647624

**Authors:** Frédéric Suffert, Kévin Meyer, Valérie Laval, Axelle Andrieux, Erik Slootweg, Pascal Frey

## Abstract

The rust fungus *Uromyces plumbarius* Peck was reported in Europe for the first time on *Oenothera lindheimeri* (Engelm. & A. Gray) W.L. Wagner & Hoch (Lindheimer’s beeblossom), a highly popular ornamental plant widely cultivated since the 2000s. The fungus was found in 11 locations across France and two locations in the Netherlands. The identification was based on an analysis of morphological characteristics and 28S rDNA sequences. The pathogenicity was confirmed through reinoculation using fresh urediniospores. This represents the first record of *U. plumbarius* outside its native range in North America. Its widespread distribution in France appears to result from rapid dissemination, likely facilitated by the plant trade between nursery gardeners. The discovery of *U. plumbarius* was accompanied by the detection of the hyperparasitic fungus *Sphaerellopsis* sp. and fungivorous larvae of *Mycodiplosis* sp.—two native, non-specific natural enemies which may play a role in limiting the impact of rust infections.

## Main text

*Oenothera lindheimeri* (Engelm. & A. Gray) W.L. Wagner & Hoch (syn. *Gaura lindheimeri*), commonly known as Lindheimer’s beeblossom or Butterfly bush (Wagner et al., 2007), is a nearly shrubby herbaceous perennial plant in the family Onagraceae, growing to 70–150 cm tall, with densely clustered branched stems. The leaves are lanceolate, finely hairy, and the stems produce clusters of small flowers opening sequentially along the open terminal panicles. *Oenothera lindheimeri* is native to southern Louisiana and Texas (USA), where it grows as a perennial plant. In Europe, it is generally treated as a perennial, but in other regions it may also be grown as an annual ornamental plant. It has been intensively propagated by nursery gardeners across many countries, particularly in Europe, including France and its neighboring countries such as Spain, Portugal, Italy, Greece, the Netherlands, and the United Kingdom. This popular ornamental plant is admired for its graceful, airy form, prolonged blooming period, and resilience to drought conditions. Several cultivars have been selected for varying flower colour, from nearly pure white to darker pink. Since the 2000s *O. lindheimeri* is frequently offered for sale in many garden centers and its presence has increased significantly in urban areas, public parks and private gardens. More recently, *O. lindheimeri* has become a common garden escape in urban areas across Europe, on waste ground or levelled soil, often in places where garden waste has been deposited (GBIF.org, 2025). In the Netherlands, for instance, it has been observed in dry grasslands within the Vlissingen port area near the railway station, where it persists in small number since its initial discovery in 2009 (Rostański & Verloove, 2015). *Oenothera lindheimeri* was also introduced in Australia some years ago and was declared an emerging, noxious weed after it escaped from cultivation.

A total of 18 samples of *O. lindheimeri* exhibiting rust were collected between 2020 and 2024 from different locations in France and in the Netherlands (Table 1; Figure 1 and 2). The infected leaves displayed uredinia on their abaxial surfaces, with several samples also showing reddish spots on the adaxial sides. Most of the uredinia, 1-2 mm in diameter, were amphigenous, scattered, at first covered by the epidermis and of a peculiar shining leaden hue, sometimes turning a darker metallic blackish brown (Figure 3). These observations align with Peck’s (1879) description of the “beautiful metallic hue of the epidermis covering the uredinia”. On young leaves—at the end of autumn on newly formed stems near the ground—uredinia initially appear as small orange blisters, 0.5–1 mm in diameter, that turn light brown and quickly rupture, exposing a mass of powdery urediniospores while remnants of the epidermis remain visible along the edges (Figure 3). On older leaves, the uredinia transition to a red-brown color and are surrounded by a reddish halo caused by the pigmentation of the plant cells. A few uredinia also develop on the stems, accompanied by reddish spots.

**Figure 1.**
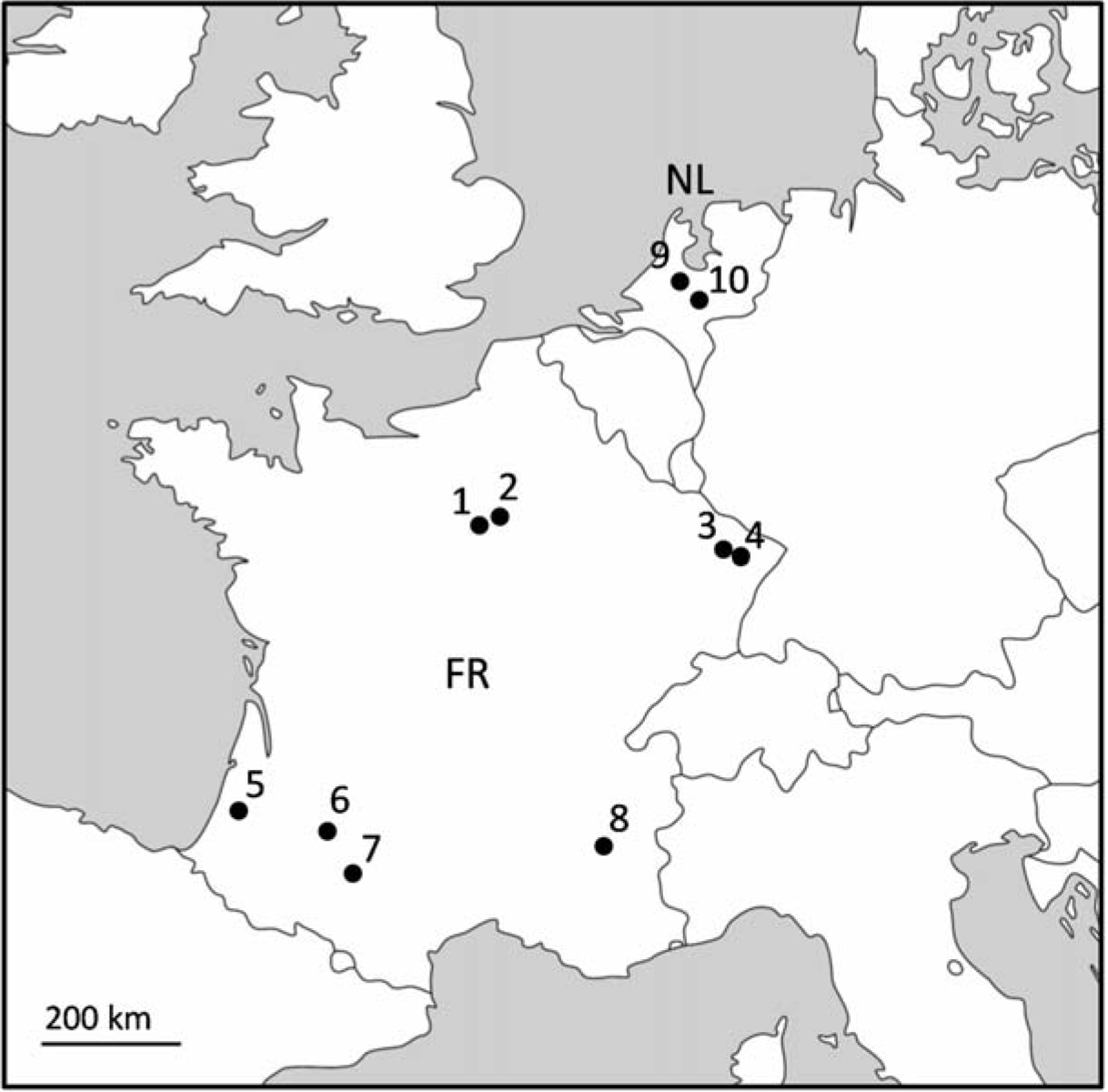
Localities in France (FR; 1. Les Clayes-sous-Bois; 2. Vincennes; 3. Nancy and Champenoux; 4. Lunéville; 5. Castets; 6. Agen; 7. Portet-sur-Garonne, 8. Villard-de-Lans) and in The Netherlands (NL; 9. Utrecht; 10. Wageningen) where *Oenothera lindheimeri* samples infected by *Uromyces plumbarius* were collected from 2020 to 2024.

**Figure 2.**
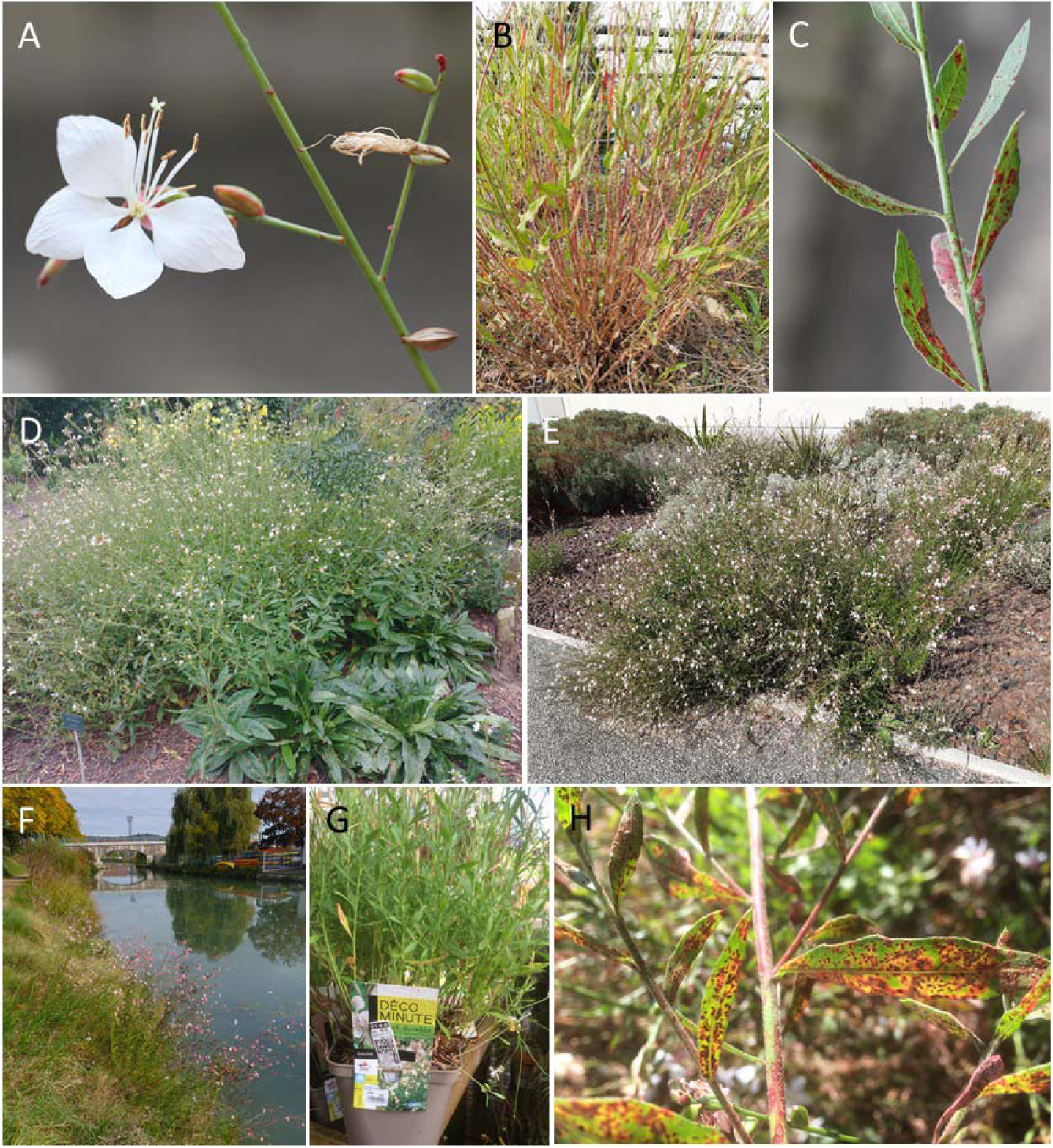
*Oenothera lindheimeri* infected by *Uromyces plumbarius.* **A-C.** Grassy embankment on the edge of a road, Les Clayes-sous-Bois, FR, 26-09-2021. **D.** Garden beds in a public park, Utrecht, NL, 17-09-2021. **E.** Garden beds of a shopping centre area, Portet-sur-Garonne, FR, 11-07-2021. **F.** Along the Canal du Midi, Agen, FR, 24-10-2021. **G.** Pots in a garden center, Nancy, FR, 09-09-2021. **H.** Garden beds, Lunéville, FR, 18-09-2021.

**Figure 3.**
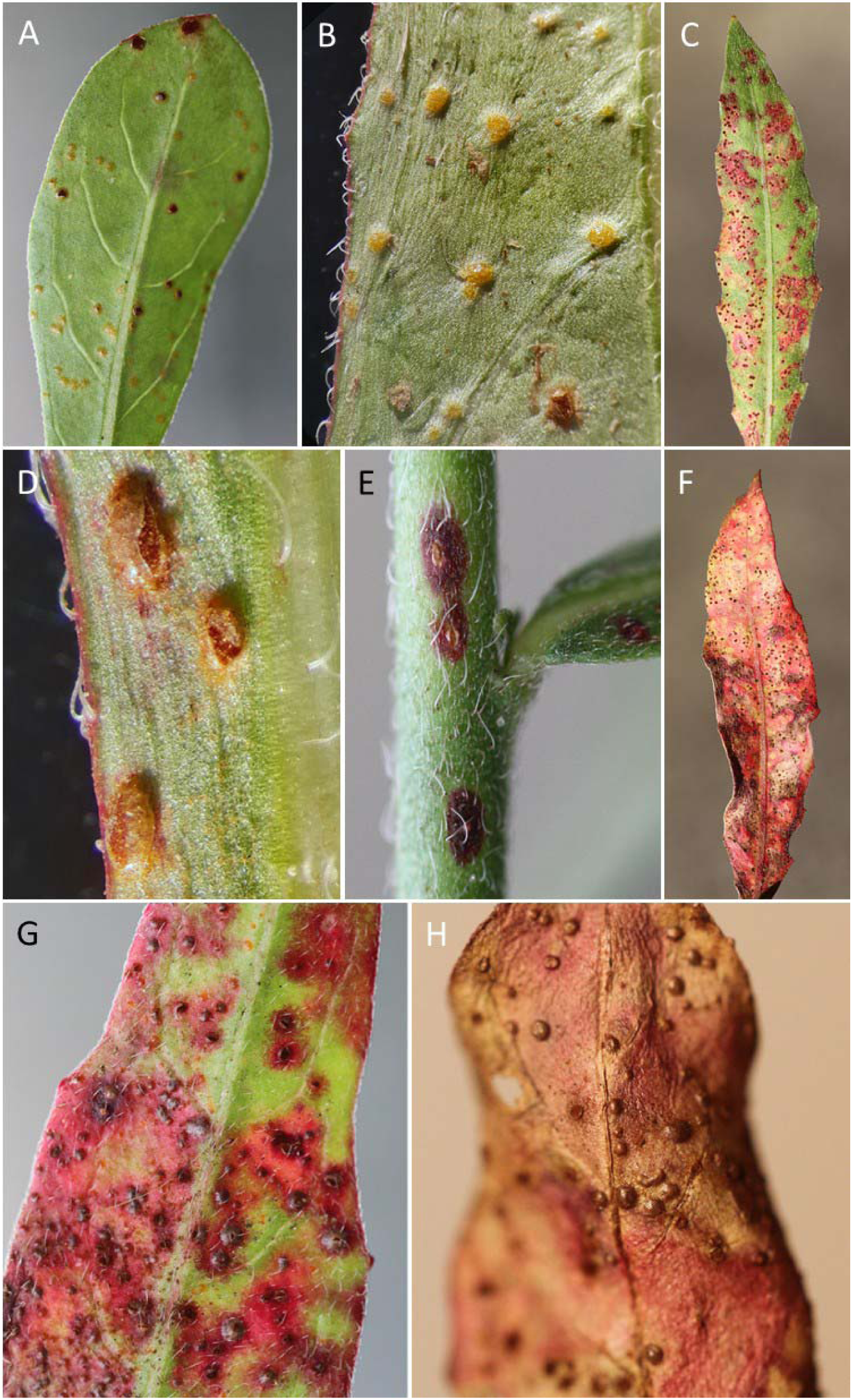
*Uromyces plumbarius* on *Oenothera lindheimeri* (Les Clayes-sous-Bois, FR). **A-B.** Immature light orange uredinia on the abaxial face of young leaves. **C.** Mature red uredinia surrounded by reddish spots on the abaxial face of leaves. **D.** Broken off uredinial epidermis revealing a mass of powdery urediniospores. E. Uredinia on a stem. **F-H.** Uredinia still protected under epidermis turning a darker metallic hue on old leaves.

**Table 1.** Origin of *Oenothera lindheimeri* samples infected by *Uromyces plumbarius* collected from 2020 to 2024 in France (FR) and in The Netherlands (NL).

| Origin | Ornamental plant status | Observer |
| --- | --- | --- |
| Portet-sur-Garonne, FR<br>11-07-2021 | garden beds in a shopping centre area | Suffert, 2021 |
| Agen, FR<br>24-10-2021 | garden beds in an urban area | Suffert, 2021 |
| Villard-de-Lans, FR<br>23-09-2021 | garden beds in an urban area | Suffert & Marcel,<br>2021 |
| Les Clayes-sous-Bois, FR<br>20-11-2020 | garden beds in an urban area | Suffert, 2020 |
| Les Clayes-sous-Bois, FR<br>26-09-2021*; 25-11-2022; 14-12-2024 | grassy embankment on the edge of a<br>road in an urban area | Suffert, 2021,<br>2022, 2024 |
| Nancy and Champenoux, FR<br>09-09-2021 | pots offered for sale in garden centers,<br>originating from two gardener nurseries<br>in Western France | Frey, 2021 |
| Lunéville, FR<br>18-09-2021; 17-10-2021 | garden beds in an urban area | Frey, 2021 |
| Vincennes, FR<br>26-09-2021 | garden beds in a public park | Marcel, 2021 |
| Castets, FR, 29-09-2021 | garden beds in an urban area | Frey, 2021 |
| Le Vésinet, FR<br>19-11-2022 | garden beds in an urban area | Suffert, 2022 |
| Wageningen, NL<br>28-08-2021; 06-11-2021 | garden beds in a public park | Slootweg, 2021 |
| Utrecht, NL<br>17-09-2021; 18-10-2022 | garden beds in a public park | Verhoeven &<br>Slootweg, 2021,<br>2022 |

\* specimen deposited in the fungal herbarium of the Muséum National d'Histoire Naturelle (Paris, France) under voucher number PC0798793

Urediniospores, present in all of the samples, are globose to slightly elongated, finely echinulate, and measure 19–24 µm by 16–19 µm on average (Table 2). Their walls are 1.5–2.0 µm thick, and they possess a short pedicel that is often absent, with two germ pores arranged supraequatorially (Figure 4). In some samples, a mixture of hyaline immature and darker mature urediniospores was observed.

**Figure 4.**
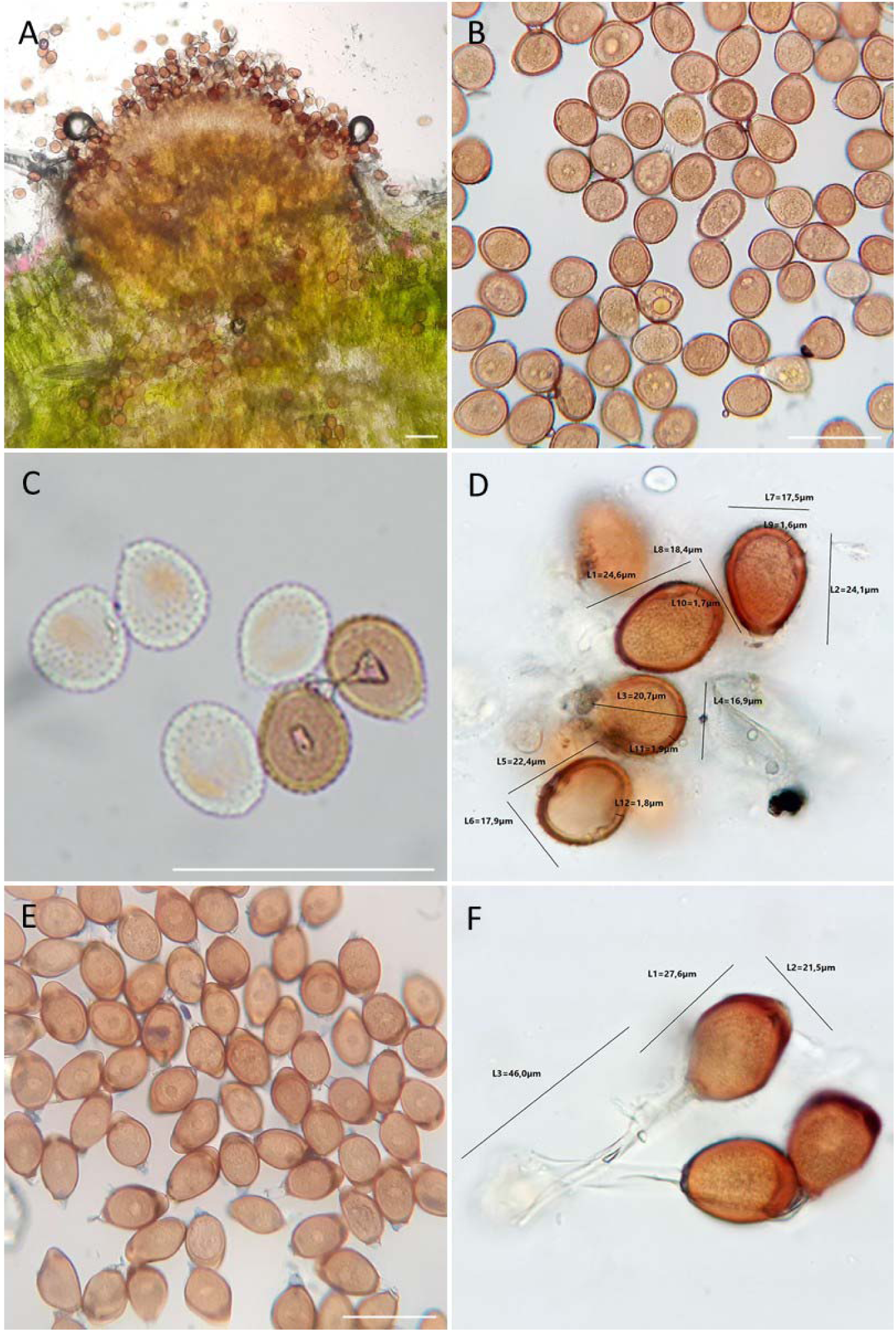
Urediniospores and teliospores of *Uromyces plumbarius.* **A.** Uredinia section. **B-D.** Urediniospores. **E-F.** Teliospores. White bars indicate 50 μm.

**Table 2.** Characteristics of *Uromyces plumbarius* urediniospores and teliospores. Measurements were carried out on ten spores of each type for every sample.

| Stage | Length | Width | Sample origin | Observer |
| --- | --- | --- | --- | --- |
| Urediniospore | 19-24 µm | 16-19 µm | Les Clayes-sous-Bois, FR<br>25-09-2021 | Suffert, 2021 |
|  | 20-24 µm | 17-19 µm | Wageningen, NL<br>28-08-2021 | Slootweg, 2021 |
|  | 22-25 µm | 16-19 µm | Utrecht, NL<br>18-10-2022 | Verhoeven &<br>Slootweg, 2022 |
| Teliospore | 24-28 µm | 17-21 µm | Les Clayes-sous-Bois, FR<br>20-11-2020 | Suffert, 2020 |
|  | 25-28 µm | 18-22 µm | Wageningen, NL<br>06-11-2021 | Slootweg, 2021 |

Teliospores were found in the first sample from Les Clayes-sous-Bois in mid-November 2020 (from *O. lindheimeri*, which was mistakenly named *Oenothera biennis* in Suffert & Suffert, 2021), in Lunéville in mid-October 2021, and in Wageningen in mid-November 2021. Teliospores are one-celled, elliptical, inconspicuously echinulate, and measure 24-28 µm x 17-22 µm on average (Table 2). Their walls are 1.0-1.5 µm thick, and they possess a long (40-45 µm) pedicel and a thickened (4-6 µm) cell wall at the apex, with one germ pore arranged mostly equatorially (Figure 4).

This description of symptoms and spore morphology aligns with previous reports on *U. plumbarius* (Peck, 1879; Saccardo, 1888; Sydow & Sydow, 1904; Bisby, 1916; Arthur & Cummins, 1962; Blomquist et al., 2015). Given the current state of knowledge, since the aecial form has never been observed on *Oenothera* sp. or any other host, *U. plumbarius* should be considered an autoecious hemicyclic rust. In our samples, uredinia and telia were indistinguishable, leaving it uncertain whether urediniospores and teliospores are produced within the same uredinia, possibly at different times. This question has not been explored in previous studies, and in retrospect, it remains unclear whether the uredinia described by Peck (1879) were considered telia, uredinia, or both.

The first leaf specimen collected in Clayes-sous-Bois (26-09-2021), exhibiting uredinia, was deposited in the fungal herbarium of the Muséum National d’Histoire Naturelle (Paris, France) under voucher number PC0798793. Some of the other samples collected in France are kept in the private herbarium of the first author. From the first leaf specimen, two samples were taken: one consisting of the entire content of an excised uredinium and the other containing only a few spores from a nearby uredinium. The DNA of both samples was extracted at INRAE BIOGER (Palaiseau) using the DNeasy Plant Mini Kit, following the manufacturer’s instructions (Qiagen). The rust fungus was identified using two pairs of rust-specific primers as follows, confirming it as *U. plumbarius* Peck. A region of the internal transcribed spacer region 1 (ITS-1) was amplified with primers ITS1rustF10d (5′-TGAACCTGCAGAAGGATCATTA-3′; Barnes & Szabo, 2007) and ITS1rustR3c (5′-TGAGAGCCTAGAGATCCATTGTTA-3′; Barnes & Szabo, 2007). A region of the ribosomal repeat spanning the 5.8S subunit, the internal transcribed spacer region 2 (ITS2), and the large subunit (28S), was amplified with primer Rust2inv (5′-GATGAAGAACACAGTGAAA-3′; Aime, 2006) and LR6 (5′-CGCCAGTTCTGCTTACC-3′; Vilgalys & Hester, 1990). PCR were performed in a 25 µl reaction volume containing DNA from each sample, 0.5 µM of each primer, 1.5 mM MgCl2, 0.2 mM of each dNTP, buffer 1X and 2 U of GoTaq G2 Flexi DNA polymerase (PROMEGA, USA). Thermocycling parameters were set as follows: 4 min at 95°C followed by 35 cycles of 30 sec at 95°C, 30 sec annealing at 56°C, 1 min at 72°C, and a final extension step of 5 min à 72°C. PCR products were visualized on a 1 % agarose gel. Successfully amplified nuclear rDNA fragments obtained were forward and reverse sequenced (Sanger Sequencing Service, Eurofins, France) using each PCR primer pairs. Forward and reverse sequences were assembled using Unipro UGENE software v51.0.0.0, followed by alignment and trimming. Sequence alignments of each ITS1 and 5.8S-ITS2-28S region showed 100% identity between the two samples, which was expected due to their close origin and the clonal reproduction typical of rust fungi. A consensus sequence was subsequently generated for each region and compared using the Basic Local Alignment Search Tool (BLASTN) to find similarities in GenBank (http://www.ncbi.nlm.nih.gov/). No match was found for the 168-bp ITS1 sequence, deposited in GenBank under accession no. PV151527 (Supplementary Material 1A) due to the absence of a reference sequence. The 1097-bp 5.8S-ITS2-28S sequence (PV151527; Supplementary Material 1B) showed 100% identity with four sequences from *U. plumbarius* specimens collected from *O. lindheimeri* in the USA: three from Louisiana (JQ312670, JQ312671, and JQ312672; Kaur et al., 2012) and one from California (KP313731; Blomquist et al., 2016). The sequence also exhibited a close relationship, with 99.8% identity to sequence GU058019, from a rust fungus sample found on *Oenothera fruticosa*, which was identified by Dixon et al. (2010) as *Puccinia dioicae*. Although *Puccinia* and *Uromyces* have been shown to be highly polyphyletic and to have evolved in close association (Maier et al., 2007; van der Merwe et al., 2007), it remains possible that this is a case of morphological misidentification, perpetuated by Kaur et al. (2012). Indeed, the identity to sequences of other specimens of this rust species available in GenBank is significantly lower, ranging from 94.0% for EF635897 to 98.4% for MK518754. The USDA fungus-host dataset of the US National Fungus Collections—the largest fungal herbarium in the Western Hemisphere— refers to 18 specimens of *P. dioicae* collected from eight different *Oenothera* species (*O. hookeri*, *O. biennis*, *O. strigosa*, *O. laciniata*, *O. fruticosa*, *O. caespitosa*, *O. mendocina*, and *O. mollisima*) in the USA, Canada, and Argentina between 1958 and 1989 (Farr et al., 2021).

However, these identifications have not been verified using molecular methods. Uncertainty remains, as *P. dioicae* is a well-established heteroecious rust species with aecial hosts in the Asteraceae and uredinial/telial hosts in the Cyperaceae. This is supported by the fungus-host dataset of the USDA fungus-host dataset, where the vast majority of the 615 recorded specimens are primarily associated with *Aster* sp., *Solidago* sp., and *Carex* sp. (Farr et al., 2021). Finally, it is worth noting that the use of the fungal-specific primers ITS1F (5’-CTTGGTCATTTAGAGGAAGTAA-3’; Gardes and Bruns, 1993) and ITS4 (5’-TCCTCCGCTTATTGATATGC-3’; White et al., 1990) on DNA extracted from the same two samples from Les Clayes-sous-Bois was unsuccessful. Moreover, amplifications performed on extracts from other leaf fragments covered with several uredinia (a few mm²) using the primer pairs Rust2inv / LR6 and ITS1rustF10d / ITS1rustR3c also failed, likely due to the presence of PCR inhibitors.

Pathogenicity tests were conducted at INRAE IAM (Champenoux) by inoculating detached *O. lindheimeri* leaves with a fresh urediniospore suspension derived from a symptomatic potted plant purchased in 2021 at the Nancy garden center (Table 1), following an inoculation protocol routinely used for poplar rusts and other rust fungi (Gérard et al., 2006). A 10-μl droplet of water agar (0.1 g.l^-1^) was deposited onto each *U. plumbarius* uredinium with a micropipette and urediniospores were dispersed within the droplet with the micropipette. The resulting urediniospore suspension was applied as 1-μl droplets on the abaxial surface of fresh leaves of *O. lindheimeri*. The inoculated leaves were incubated on wet filter paper in 24 x 24 cm Petri dishes, at 19±1°C under continuous LED illumination (50 mmol.s^-1^.m^-2^) in an incubation room. New uredinia appeared between 12 and 20 days post-inoculation.

Fungal specimens of the genus *Sphaerellopsis* (sexual morph *Eudarluca*), known to include several cosmopolitan hyperparasitic species occurring on a wide range of rust fungi (Trakunyingcharoen et al., 2014), were observed in two *U. plumbarius* samples: one associated with uredinia and telia collected from Wageningen (06-11-2021) and the other associated with uredinia from Les Clayes-sous-Bois (27-11-2021) (Figure 5). The conidiomata of *Sphaerellopsis* sp., which pierced the uredinia, were globose, measuring 150–250 µm in diameter, dark brown, and featured a central ostiole exuding numerous conidia aggregated into a white, filamentous cirrhus of varying thickness. The conidia were hyaline, 1-septate, fusiform, measuring 13.5–17.5 μm × 3.5–4.5 μm, slightly constricted at the septa, and possessed funnel-shaped mucoid appendages at both ends. To our knowledge, this is the first report of *Sphaerellopsis* sp. infecting *U. plumbarius*. Our observations align with the description by Trakunyingcharoen et al. (2014) of *Sphaerellopsis filum*, which has been recorded in association with 369 rust species from 30 genera across more than 50 countries (Kranz & Brandenburger, 1981). Nonetheless, the recently revealed diversity within the genus *Sphaerellopsis* (Gómez-Zapata et al., 2024a) suggests that many historical identifications of *S. filum* likely encompassed cryptic or unrelated taxa, and that reliable species delimitation based solely on morphology is problematic. In the absence of molecular analyses—given that the specimen isolated from Les Clayes-sous-Bois was not preserved—it was impossible to confirm the species identity.

**Figure 5.**
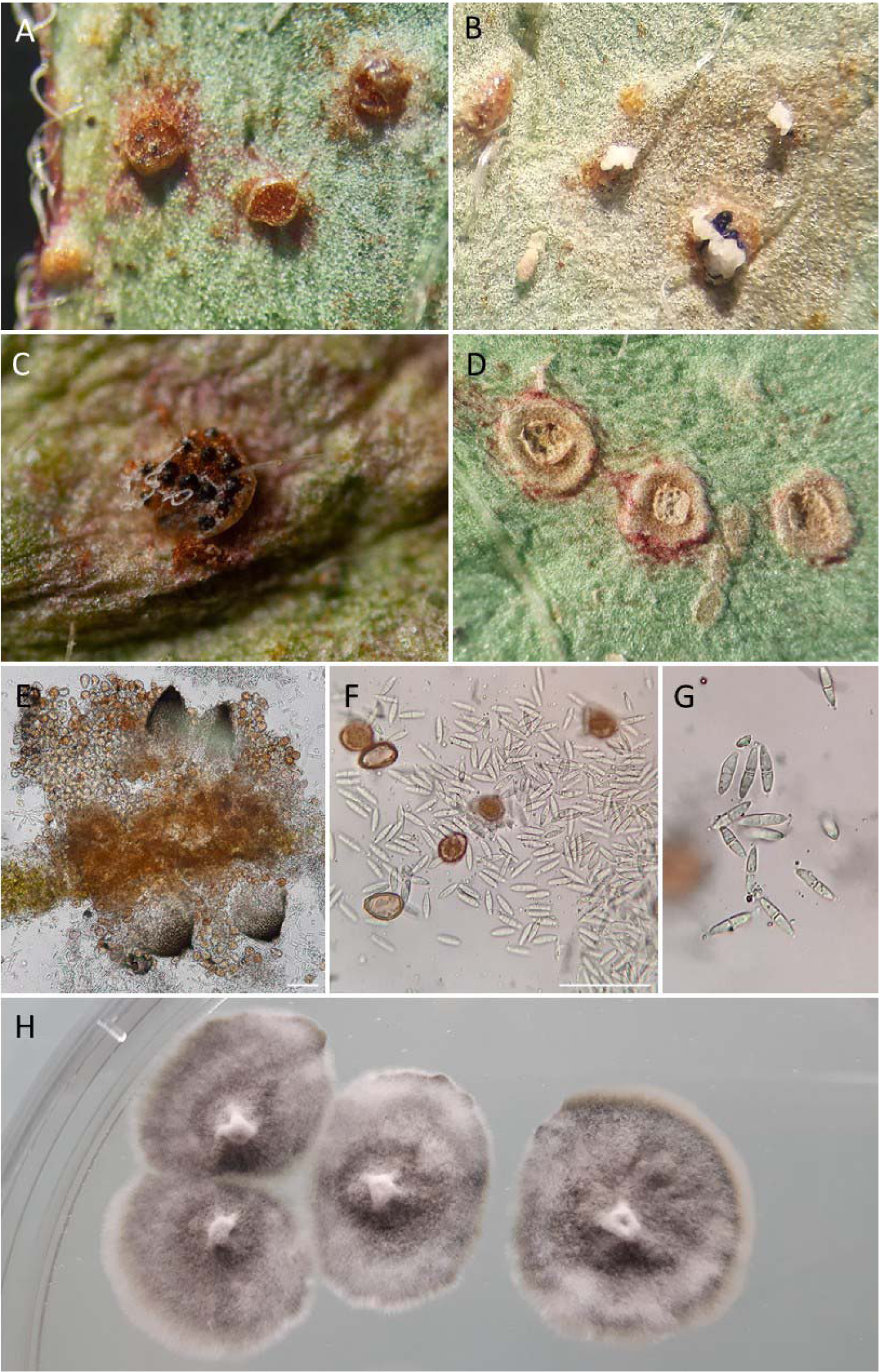
*Sphaerellopsis* sp. parasiting *Uromyces plumbarius* on *Oenothera lindheimeri.* **A.** Mature *U. plumbarius* uredinia parasitized by *Sphaerellopsis* sp., with one pierced by three conidiomata. **B-C.** Cirrhi (white filaments) extruded from conidiomata (black). **D.** Insertion scars from old *U. plumbarius* uredinia and *Sphaerellopsis* sp. conidiomata. **E.** Section of *U. plumbarius* uredinium and *Sphaerellopsis* sp. conidiomata. **F-G.** *Sphaerellopsis* sp. conidia. **H.** Colonies of *Sphaerellopsis* sp. growing on PDA medium in a Petri dish.

**Figure 6.**
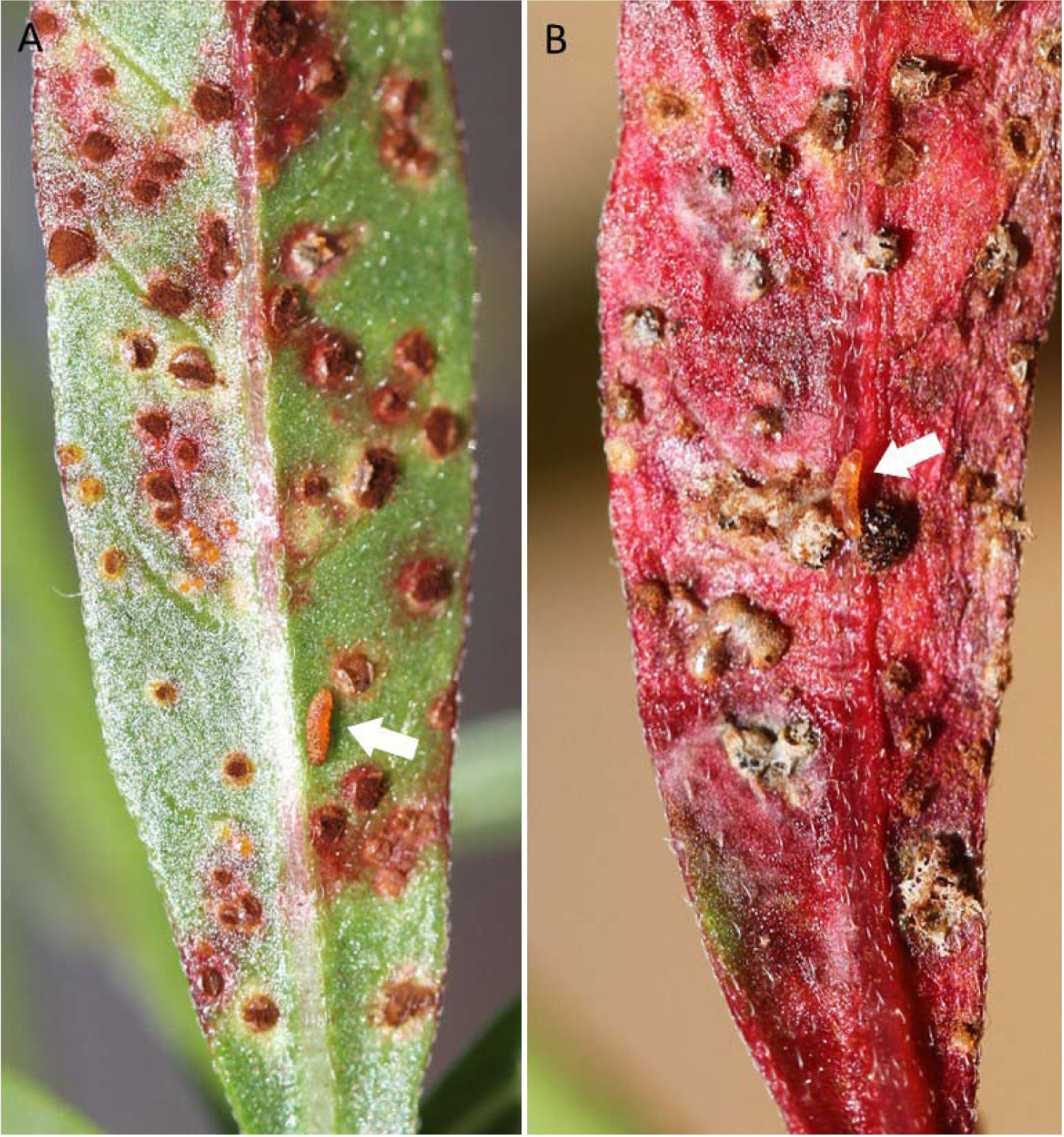
**A-B.** Larvae of *Mycodiplosis* sp. (white arrow) feeding on (**A**) fresh uredinia of *Uromyces plumbarius* on *Oenothera lindheimeri* and (**B**) older uredinia heavily parasitized by *Sphaerellopsis* sp.

Gall midge fly larvae of the genus *Mycodiplosis*—another well-known natural enemy of rust (Henk et al., 2011)—were observed in two *U. plumbarius* samples: one from Les=Clayes-sous-Bois (26=September=2021) and the other from Agen (24=October=2021) (Figure=6). Although morphological characters readily allowed identification to the genus level, which comprises 49 species (Gagné and Jaschhof 2021, Kolesik et al. 2022), species-level determination was not conducted.

This report documents the presence of *U. plumbarius* across a dozen locations, spanning more than 1,500 km between the most distant sites, with symptomatic plants consistently observed. To our knowledge, *U. plumbarius* has not been previously recorded outside the USA and Mexico GBIF.org, 2024), making this its first report in Europe. Furthermore, this constitutes the first detection of a rust fungus on the genus *Oenothera* in Europe. These preliminary data suggest that *U. plumbarius* might be widely distributed across Western Europe, wherever *O. lindheimeri* is cultivated, therefore considered newly endemic to the area.

It can be assumed that the pathogen was introduced from North America with one of its host plants—most likely *O. lindheimeri*—and subsequently spread through the propagation and distribution of plants by gardener nurseries and garden centers. This hypothesis of a recent introduction aligns with the fact that no rust species has yet been described on this genus in Europe (Klenke & Scholler, 2015). According to Wagner et al. (2007) the centre of diversity of Onagraceae is in southwestern North America, and the family is now widely distributed in temperate to subtropical regions of the Americas (Rostański et al., 1994; Rostański & Verloove, 2015). Several *Oenothera* species have become naturalized nearly worldwide, most likely in post-Columbian times. Many of these species are now found in environments heavily disturbed by human activity, such as railway grounds, gravel pits, dredge dumping areas, sand and mineral depots, and ruderalized sand dunes. Whether several species are endemic to or originated in Europe is still debated. The consensus, however, is that *O. biennis* has been naturalized in Europe—most likely in the Western European region encompassing Belgium, northern France, and the Netherlands—since the late 18th century (Rostański & Verloove, 2015; Roucel, 1792). Many other taxa are considered either newcomers that arrived during the 19th century or neotaxa that evolved from the original species.

The synthesis of *U. plumbarius* records in the GBIF.org database GBIF.org, 2024), reports 391 occurrences between 1819 and 1994, with a distribution across the entire USA and a presence in Mexico. Numerous specimens of this rust species are preserved in the fungal herbariums of university laboratories, museums and USDA. *U. plumbarius* was reported on different Onagraceae taxa by Bisby (1916), based on a literature review without any validation of individual host: *Anogra* sp., *Boiduvalia* sp., *Chamanirion* sp., *Chylisma* sp., *Circaea* sp., *Clarkia* sp., *Epilobium* sp., *Eulobus* sp., *Fuchsia* sp., *Gaura* sp., *Gayophytum* sp., *Godelia* sp., *Jussiaea* sp., *Kneiffia* sp., *Ludwigia* sp., *Luxavia* sp., *Megapterum* sp., *Meriolix* sp., *Oenothera* sp., *Onagra* sp., *Pachylopus* sp., *Phaestoma* sp., *Sphaerostigma* sp., *Taraxia* sp., *Zauchneria* sp. Concerning *Oenothera* sp., *U. plumbarius* was notably reported on *O. biennis* in Illinois (Manaaki Whenua, 2002), Ohio (Ellett, 1955) and Iowa (Tiffany & Knaphus, 2004); on *O. caespitosa* in Colorado (Peck, 1879); *O. linderheimeri* in Ohio (Ellett, 1955), Iowa (Tiffany & Knaphus, 2004), Louisiana (Kaur et al., 2012; Ferrin, 2012) and California (Blomquist et al., 2015); on *O. laciniate* and *O. simulans* (syn. *Gaura angustifolia*) in Mississippi (Presley, 1947); on *O. sinuosa* in Texas (Manaaki Whenua, 2002); on *O. suffrutescens* (syn. *Gaura coccinea*) in Nebraska (Manaaki Whenua, 2002), Texas (Manaaki Whenua, 2002), Mississippi (Presley, 1947), Colorado (Manaaki Whenua, 2002), and in Mexico (Hennen & Cummins, 1967). This list can be further supplemented by consulting the USDA fungus-host dataset of the U.S. National Fungus Collections (Farr et al., 2021).

While the impact of *U. plumbarius* on the growth of *O. lindheimeri* seems currently limited, gardeners may have to contend with this rust in the future, as it could diminish the ornamental value of the plants due to the highly visible reddish discoloration it induces on the leaves and stems. A concern is that plants sold in garden centers in France (in Nancy and Champenoux; Table 1) and in the Netherlands (Ochten; Erik Slootweg, pers. comm.) were found to be infected. It is thus therefore that gardener nurseries, along with potentially several others, have unintentionally contributed to the spread of this rust in Europe through the trade of *O. lindheimeri*. As this species is still gaining in popularity as a garden plant, *U. plumbarius* will probably remain well established in Europe. A subsequent concern to address is the risk of a potential host jump by *U. plumbarius* to other European wild *Oenothera* species (27 recorded since 1950; Dietrich et al., 1997; Rostański et al., 1994), for instance *O. biennis*, *O. erythrosepala*, *O. silesiaca*, *O. fallax*, *O. glazioviana*, *O. deflexa*, which are quite common in France and Belgium. In Europe, only few species are successful in terms of increasing abundance while the majority of the species remain rare and local, constituting a wealth in terms of biodiversity. This issue is all the more preoccupying since *U. plumbarius* was identified on some other *Oenothera* species in the USA (see above).

Of the 27 species of rust fungi on members of the Onagraceae family described worldwide at the beginning of the last century (Sydow & Sydow, 1915; Brisby, 2016), 15 were reported as occurring exclusively in North America and three only in South America. Several species are common in Europe, particularly on various plant hosts in the genera *Circaea* and *Epilobium* (Klenke & Scholler, 2015). The vast majority of these species belong to the genera *Puccinia* and *Pucciniastrum*, with two *Aecidium* forms and one *Uredo*. The only species from the genus *Uromyces* is *U. plumbarius*. Brisby (1916) notes that some authors once considered *U. plumbarius* to represent a combination of morphological races—or even four distinct species—on different host species, but this view has not stood the test of time.

The report of *Sphaerellopsis* sp. associated with *U. plumbarius* in Les Clayes-sous-Bois and Wageningen suggests that a natural balance may already be establishing between the pathogen and hyperparasite populations, potentially contributing to the limitation of rust attacks. While complete natural regulation can be ruled out, it has been suggested that several *Sphaerellopsis* species could help reduce losses caused by certain rusts (e.g., Yuan et al., 1999; Gordon & Pfender, 2012; Kajamuhan et al., 2015). This requires that the local population of the hyperparasite fungus be sufficiently present at the start of the season, enabling it to increase rapidly enough to slow rust development before the disease becomes damaging. This can occur if a population of another rust species on a weed preexists, creating a reservoir from which the hyperparasite fungus can spread to *U. plumbarius*. This was the case in Les Clayes-sous-Bois, where infected *O. lindheimeri* were found along a grassy embankment on the edge of a road, in direct contact with grasses infected by *Puccinia* sp., themselves hyperparasitized by *Sphaerellopsis* sp. The results of Kajamuhan et al. (2015) on several rusts of grasses demonstrate substantial and wide variation in naturally occurring populations of *Sphaerellopsis filum* (sexual morph *Eudarluca caricis*), with some evidence of specialization within sympatric populations. Understanding such complex patterns would be valuable in developing effective strategies for promoting natural regulation or biological control, especially considering that significant effort is still needed to clarify the identity of *Sphaerellopsis* species. Literature prior to Trakunyingcharoen et al. (2014) often conflated multiple *Sphaerellopsis* species under the name *S. filum*, a practice now recognized as problematic (Gómez-Zapata et al., 2024a).

The report of *Mycodiplosis* sp. larvae feeding on the spores of *U. plumbarius* in Les Clayes-sous-Bois and Agen (France) is also of interest, even though the species remains unidentified. Given the relatively broad host range of this insect genus—recently highlighted through analyses of phylogenetic relationships, biogeography, and host range (Gómez-Zapata et al., 2024b)—this suggests that their role in regulating populations of certain rust fungi could be significant.

To conclude, this new geographic record of a known rust fungus species, found in association with two native, non-specific natural enemies on a recently introduced ornamental plant host, represents a noteworthy epidemiological situation. Invasions by pathogenic fungi are usually characterized by an absence of coevolution between the fungus and its new hosts, referred to as “new encounter” or “novel interaction” (Parker & Gilbert, 2004; Desprez Loustau et al., 2007). This concept can be extended to the interactions between *U. plumbarius* and native natural enemies—in this case, a hyperparasitic fungus and a fungivorous insect —which were, in all likelihood, already present in Europe prior to the arrival of the rust fungus. These interactions therefore represent a second level of “new encounter.” The trophic network now formed by the links *O. lindheimeri - U. plumbarius - Sphaerellopsis* sp. *-Mycodiplosis* sp. offers a compelling illustration of what may occur when, in efforts to manage a plant disease, the presence (or even introduction) of biocontrol agents is encouraged—agents whose populations or strains are endemic, but have not previously coexisted with the invading pathogen. Investigating the host range of the hyperparasite and the feeding behavior of the fungivore, as done in parallel by Gómez-Zapata et al. (2024a, 2024b), is of practical relevance when aiming to promote these interactions. Additionally, this case holds theoretical significance for testing hypotheses concerning the dynamics and functioning of co-evolutionary pathosystems (Thompson, 1999).

## Data availability

Consensus sequences of the specimen *Uromyces plumbarius* are available online via the Gene Bank Information System at https://www.ncbi.nlm.nih.gov/genbank/ (accession no. PV153707 and PV151527).

---

**Supplementary Material 1.**
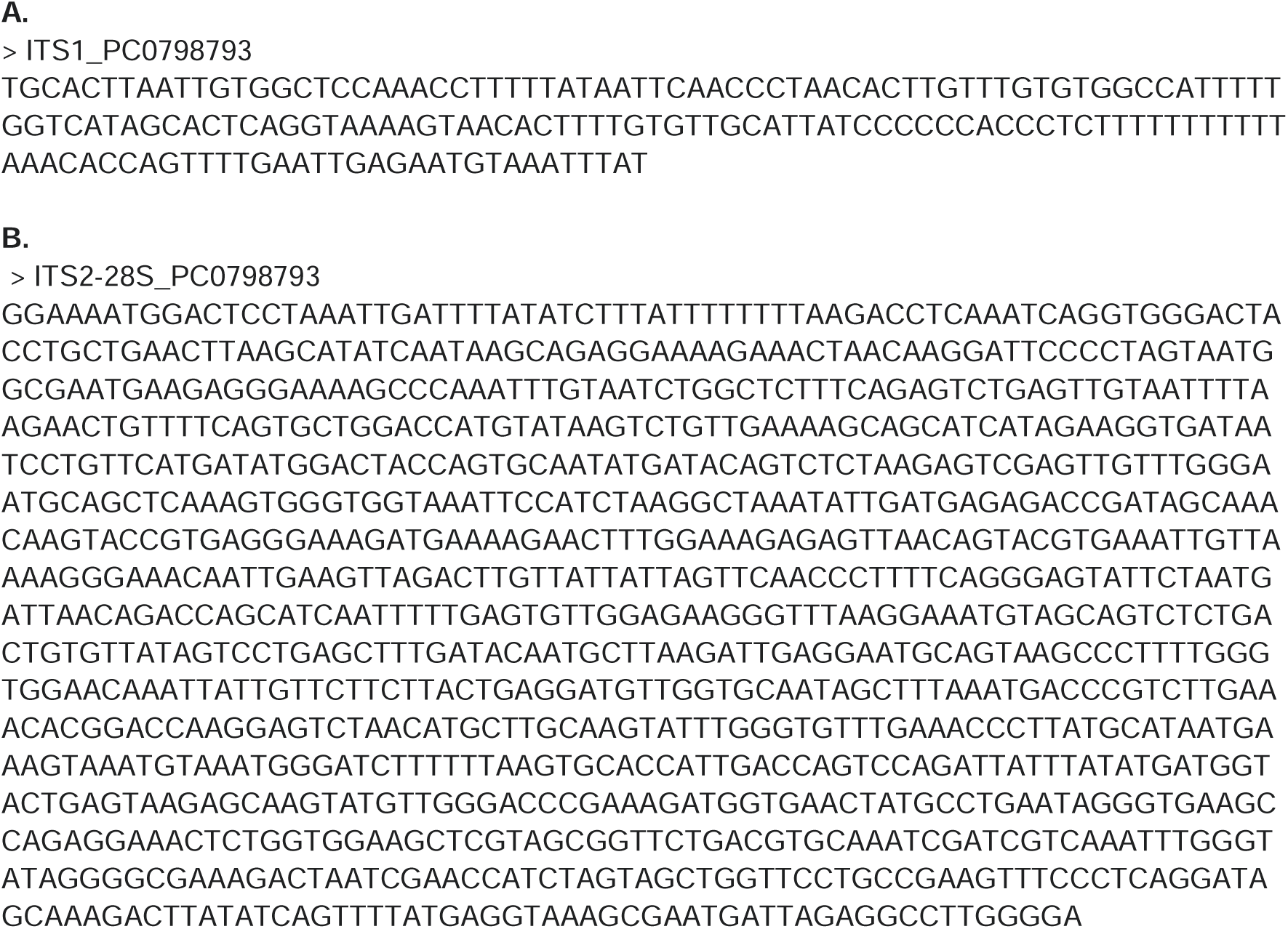
Consensus sequences of the specimen *Uromyces plumbarius* obtained with the pairs of primers (**A**) ITS1F10D / ITSRustR3C (GeneBank accession no. PV153707) and (**B**) Rust2inv / Lr6 (GeneBank accession no. PV151527)

